# Single-cell and neuronal network alterations in an *in vitro* model of Fragile X syndrome

**DOI:** 10.1101/366997

**Authors:** Anastasiya Moskalyuk, R. Frank Kooy, Michele Giugliano

## Abstract

The Fragile X mental retardation protein (FMRP) is involved in many cellular processes and it regulates synaptic and network development in neurons. Its absence is known to lead to intellectual disability, with a wide range of co-morbidities including autism. Over the past decades, FMRP research focused on abnormalities both in glutamatergic and GABAergic signalling, and an altered balance between excitation and inhibition has been hypothesised to underlie the clinical consequences of absence of the protein. Using FMRP knockout mice, we studied an *in vitro* model of cortical microcircuitry and observed that the loss of FMRP largely affected the electrophysiological correlates of network development and maturation but caused less alterations in single-cell phenotypes. Using a mathematical model, we demonstrated that the combination of an increased excitation and reduced inhibition describes best predicts our experimental observations during the *ex vivo* formation of the network connections.

## Introduction

Our cognitive abilities depend to a large extent on the normal development and functioning of the neuronal networks in the brain. Genetic abnormalities that disturb the establishment of physiological neuronal connectivity lead to a variety of disorders, of which Fragile X syndrome (FXS) is an important example (Contractor et al. 2015). FXS is a frequent cause of autism and intellectual disability and it is caused by the absence of the FMR1 protein (FMRP), mostly due to promotor hypermethylation (Santoro et al. 2012). The function of the underlying gene has been extensively studied and FMRP quickly became recognized for its pivotal role in many cellular processes, including translation, RNA transport, and stability (Penagarikano et al. 2007; Willemsen and Kooy 2017).

However, despite over 20 years of research, it still remains elusive how the loss of FMRP affects brain connectivity. Abnormalities in both excitatory and inhibitory transmission have been reported in FXS. For instance, it has been proposed that the cognitive symptoms of the disease are due to the hyperactivation of group I metabotropic glutamate receptors (mGluR) (Bear et al. 2004). This theory has been confirmed in animal studies but the results in human clinical trials failed to meet the expectations (Dolen et al. 2007; Berry-Kravis et al. 2012). In addition, a reduction in the γ-aminobutyric acid (GABA) pathway has been implicated in FXS and was hypothesised to play a role in specific aspects of its clinical presentation (D’Hulst et al. 2006; Gantois et al. 2006; D’Hulst and Kooy 2007; Braat and Kooy 2015). Indeed, measurable effects on GABAergic currents could be demonstrated in patch-clamp *in vitro* experiments (Ligsay et al. 2017; Sabanov et al. 2017).

Other studies, performed at the (sub)cellular level, have examined structural neuronal connectivity in Fmr1 KO mice while focusing on abnormalities in the morphology of the dendritic spines. A higher dendritic spine density was observed in the cortex and hippocampus of Fmr1 KO adult mice (Galvez and Greenough 2005; McKinney et al. 2005), although the opposite finding was also reported (Braun and Segal 2000). Despite such apparent inconsistencies, most studies converge in suggesting that an immature dendritic spine phenotype and an abnormal synaptic connectivity are typical features of Fmr1 KO mice (reviewed by He and Portera-Cailliau 2013). Other studies focused on synaptic transmission and reported defects in presynaptic neurotransmitter release and/or on postsynaptic receptor sensitivity.

In this study, we focused on an intermediate level of organization of the nervous system in FXS mice, increasing the complexity of existing (sub)cellular assays: the level of a generic cortical microcircuit. By studying the electrophysiological correlates of cellular and synaptic abnormalities in FXS, we aimed to link together the known defects in synaptic transmission to the network activity. We further aimed to test the hypothesis that at different stages of network development, the balance between excitation and inhibition might be different. We finally employed a minimal computational model to support our conclusions on an overall dysfunctional network organization, consisting of alterations in both synaptic excitation and synaptic inhibition.

## Materials and Methods

**Neuronal cell cultures**. *Fmr1* knockout and wild-type colonies were generated by crossing females heterozygous for the *Fmr1* mutation (B6.129P2-*Fmr1*^tm1Cgr^/Ant backcrossed for more than 20 generations to C57BL/6 J) with knockout and C57BL/6 J wild-type (Charles River, Wilmington, MA, USA) males, respectively. Genotypes were determined by PCR on DNA isolated from tail biopsies (Bakker et al. 1994). All animals were housed in groups of approximately 5 littermates in standard mouse cages under conventional laboratory conditions, i.e., food and water *ad libitum*, constant room temperature and humidity, and a 12:12 h light-dark cycle.

We employed newborn (male and female) C57BL/6J wild-type and Fmr1-KO1 mice, backcrossed on a Harlan C57BL/6J background strain for more than 20 generations, for neuronal primary cultures. We prepared cells as described previously (Pulizzi et al. 2016), euthanizing the pups by rapid decapitation and closely following the international guidelines on animal welfare. All procedures were approved by the Ethical Committee of Antwerp University (permission no. 2011_87) and licensed by the Belgian Animal, Plant and Food Directorate-General of the Federal Department of Public Health, Safety of the Food Chain and the Environment (license no. LA1100469).

We used microelectrode arrays integrated into glass substrates (Fig. 1) (MEAs; MultiChannel Systems, Reutlingen, Germany) as well as conventional glass coverslips for network-level electrophysiology and single-cell patch-clamp experiments, respectively. Prior to cell seeding, we treated their surface with polyethyleneimine (PEI, 0.1% wt/vol in milli-Q water at room temperature, Sigma-Aldrich, Germany) and then rinsed with milli-Q water and air-dried it. We seeded cells with an initial density of 6’500/mm^2^ on MEAs, and 1’500 cell/mm^2^ on coverslips. After seeding, we maintained cells in a conventional incubator at 5% CO_2_, 37°C, and 95% humidity (5215, Shellab, Cornelius, OR, USA). During culturing and electrophysiological recordings, we sealed MEAs with fluorinated Teflon membranes (Ala-MEA-Mem, Ala Science, Farmingdale, NY, USA), reducing risks of contamination, preventing water evaporation and alteration of osmolarity, and ensuring O_2_ and CO_2_ gas exchanges. After 8 days *in vitro* (DIV8), we added fresh and pre-warmed medium to reach a 1 ml final volume. From DIV10 onwards, we replaced half of the culture medium volume with fresh, pre-warmed medium, every 2 days. We obtained all reagents from Sigma-Aldrich (St. Louis, MO, USA) or Life Technologies (Ghent, Belgium).

**Figure 1.**
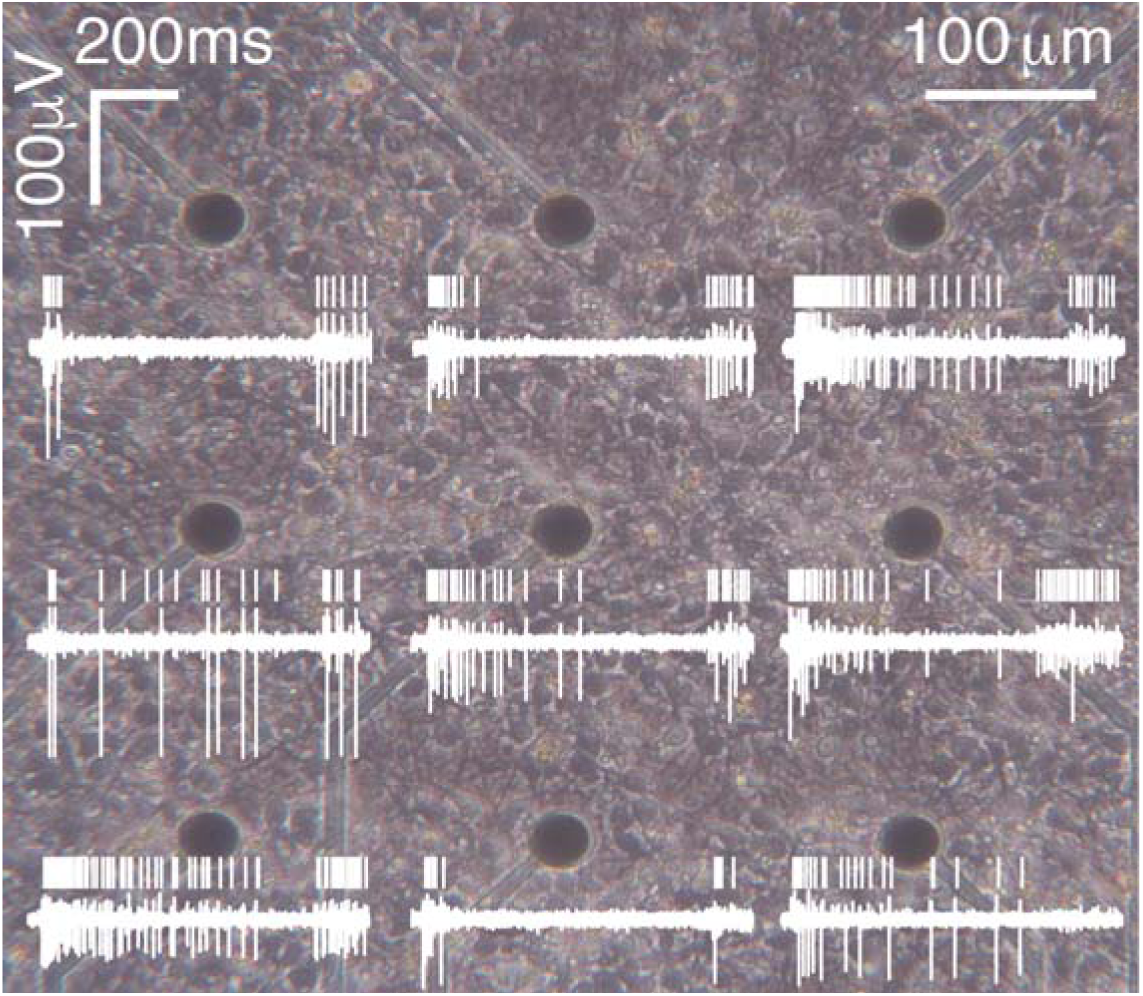
Cortical neurons, dissociated from the newborn murine cortex, form electrically active networks *in vitro*. The 20x phase-contrast micrograph shows cells 21 days after plating on the surface of an 8 by 8 array of microelectrodes (MEA), integrated in a glass-substrate. Nine out of 60 microelectrodes are visible as darker circles, with 30 μm diameter and 200 μm spacing. The white traces represent 800 ms long high-passed recordings of the extracellular electrical potential of the corresponding microelectrodes. The computed times of occurrence of action potentials (AP) are depicted above the raw traces, as detected by an adaptive signal-processing algorithm. Note the degree of AP synchronization across the nine microelectrodes.

**Network electrophysiology: experiments and data analysis**. We employed commercial MEAs, whose 60 titanium nitrate (TiN) microelectrodes were arranged in an 8×8 regular layout (60MEA200/30iR-ITO-gr, MultiChannel Systems, Reutlingen, Germany). Each microelectrode had a 30 μm diameter and a 200 μm spacing from its nearest neighbors and was used for the non-invasive long-term monitoring of the spontaneous electrical activity of cells, throughout network development *ex vivo*. We followed each MEA for up to 35 days *in vitro* (DIVs), performing each recording for at least 30 minutes, at 37°C and under 5% CO_2_. We detected extracellular raw electrical potentials at each of the 60 microelectrodes of an MEA and amplified them by a MEA-1060-Up-BC electronic multichannel amplifier, with a 1–3000 Hz bandwidth and an amplification factor of 1200 (Multichannel Systems, Germany). We sampled raw analog signals at 25 kHz/channel and digitized them at 16 bits by an A/D electronic board (MCCard, MultiChannel Systems). We finally stored data on disk by means of the MCRack software (Multi Channel Systems) for subsequent analyses. All data processing was performed off-line, using custom-written MATLAB scripts (The MathWorks, Natik, MA, USA). Action potentials (APs) and network-*burst* detection was carried out using QSpike Tools (Mahmud et al. 2014). Briefly, we determined the times of occurrence of AP by an adaptive peak-detection algorithm, following a band-pass filter (300-3000 Hz) of the raw electrical potential waveform recorded from each microelectrode. This algorithm identified as a putative AP each threshold-crossing event, exceeding five times the standard deviation of the background noise (Quiroga et al. 2004).

We identified network-*burst* as major synchronization events, each containing AP in at least 10% of active microelectrodes in a 1 ms time window, with an artificial refractory period for their detection of 50 ms. The on-and offsets of a *burst* were defined as the times around the *burst* peak activity, where the Gaussian-smoothed network-wide spike-time histogram (STH, estimated using bins of 1 ms) reached zero. We derived basic statistics on *burst* occurrence frequency, duration and inter-burst intervals from these events. We further examined the time course of the firing rate during each *burst* (i.e. STH), in terms of spectral (dominant) frequency content following the burst peak, if present. We revealed the oscillatory frequency content in the offset phase of each *burst*, by pre-processing the data and individually aligning each burst prior to averaging, maximizing similarity, as described previously (Pulizzi et al. 2016). We performed frequency-domain analysis to extract the “dominant” frequency of the oscillation, by estimating the spectrogram of each STH. In detail, we employed the Fast Fourier Transform algorithm to extract the time-varying spectrum of frequencies contained in the STH, averaging over all *bursts*. We defined as the “dominant” component of the power spectrum the frequency corresponding to the highest peak in the spectrum, which exceeded the median value of the spectrum by at least three-fold.

**Single-cell electrophysiology: experiments and data analyses**. For investigating single-cell excitability and synaptic currents, we performed patch-clamp experiments in the whole-cell configuration from the soma of individual neurons. We carried out both current- and voltage-clamp recordings by an Axon Multiclamp 700B amplifier (Molecular Devices LLC, US). Current and voltage traces were low-pass filtered at 3 kHz, sampled at 20 kHz, digitized at 16 bits by a National Instruments A/D board, and stored for offline analyses by the software LCG (Linaro et al. 2014). We then processed the data with custom scripts, written in MATLAB (The MathWorks, Natick, US).

We pulled patch electrodes from standard borosilicate glass capillaries (1BF150, World Precision Instruments, UK), by means of a horizontal puller (P97, Sutter, Novato, US), with a resistance of 5.5-7.5 MΩ when filled with an intracellular solution containing (in mM): 135 K-gluconate, 10 KCl, 10 HEPES, 4 Mg-ATP, 0.3 Na_3_GTP (pH 7.3, adjusted with KOH). We obtained all recordings at 34 °C and upon replacing the culture medium by an extracellular solution, constantly perfused at a rate of 1ml/min, and containing (in mM): 145 NaCl, 4 KCl, 2 Na-pyruvate, 5 Hepes, 5 glucose, 2 CaCl_2_, and 1 MgCl_2_ (pH adjusted to 7.4 with NaOH). For isolating spontaneous excitatory (sEPSC) and inhibitory (sIPSC) currents, we used the voltage-clamp mode and held neurons for 5 min at a holding potential of −70mV or of 0mV, thus matching the inhibitory (Cl^-^) or the excitatory (K^+^, Na^+^) Nernst reversal potentials, respectively.

**Statistics**. We verified the normality of the distribution of each acquired parameter by the Lilliefors test. For normal distributions, we showed data as mean ± standard error of the mean (SEM), and we assessed the statistical significance of differences between groups by a two-way ANOVA and post-hoc Fisher's procedure (Fig. 2), t-test (Figs. 6-7). In case of non-normal distributions, we used the median and the interquartile range for data presentation, and we assessed the statistical significance of differences between groups by a Mann-Whitney non-parametric test for two unpaired groups (Fig. 3). Differences with a value of p < 0.05 (*), p < 0.01 (**) and of p < 0.001 (***) were considered significant.

**Figure 2.**
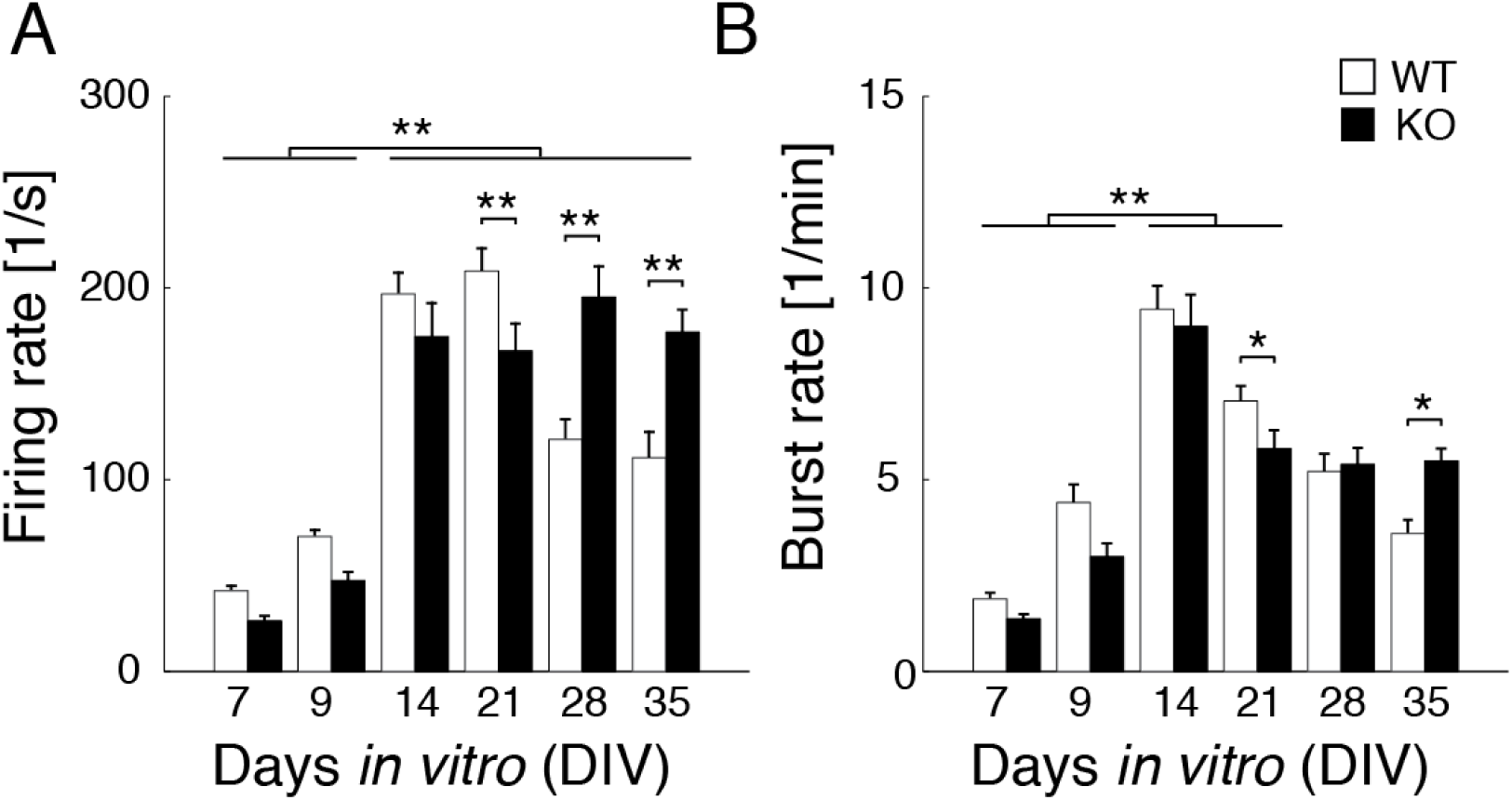
Coarse analyses of the network activity during development *ex vivo*. *In vitro* neuronal networks display spontaneous episodic synchronization of APs, here named *bursting* (Fig. 1). The mean rate of firing (**A**) recapitulates, in both Fmr1 KO and WT networks, a well-known progression over five weeks of *in vitro* development: the MEA microelectrodes detect significantly less events at DIV7-9 than at older stages. At DIV21, KO (black bars) and WT (white bars) significantly differ (p < 0.01). While WT show a characteristic decrease in the mean firing rate after DIV21, KO networks retain a higher rate that was significantly different than controls. Comparable observations are obtained when the mean rate of *burst* synchronization (**B**) is quantified. Both KO and WT networks generate more *bursts* at DIV14-21 than at DIV7-9 (p < 0.01). However, while WT display a characteristic decrease in the mean *burst* rate after DIV14, KO shows a persisting higher mean *bursting* rate at full *ex vivo* maturity (DIV35; p < 0.05). Data are displayed as mean ± SEM.

**Mathematical model**. We employed a minimal mathematical model of a neuronal network (Wilson and Cowan 1972; Amit and Tsodyks 1991; Dayan and Abbott 2001; Pampaloni et al. 2018) to describe the mean firing rates *ν_E_(t)* and *ν_I_(t)* of excitatory and inhibitory neurons, reciprocally and recurrently connected. Following previous work (Giugliano et al. 2008; La Camera et al. 2008; Gambazzi et al. 2010; Gigante et al. 2015), we associated with each neuronal subpopulation a characteristic time scale (i.e. *τ_E_* and *τ_I_*) as well as a single-cell *f-I* curve (i.e. *ϕ*(*I*)).

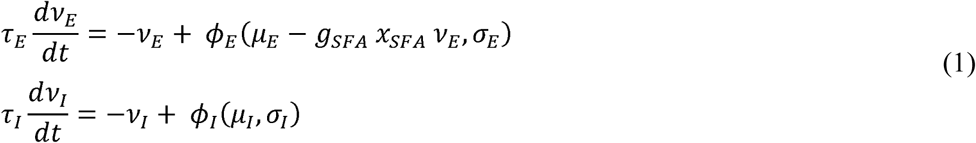

Excitatory neurons included a negative-feedback from spike-frequency adaptation mechanisms, by a slow variable *x_SFA_* evolving as

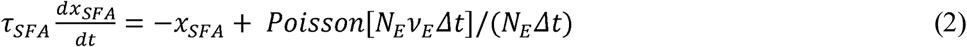

where *N_E_* is the number of excitatory neurons, *τ_SFA_* the adaptation time scale, Δ*τ* the simulation time step, and Poisson[m] is a random variable of mean *m*, capturing finite-size effects. The synaptic input to each neuron was specified in terms of (infinitesimal) mean *μ* and variance *σ*^2^, under the hypotheses of the extended mean-field theory (La Camera et al. 2008). These reflected external inputs and the synaptic connectivity (Fig. 8A), through the size of presynaptic populations (i.e. *N_ext_, N_E_, N_I_*), the probability of recurrent connectivity (i.e. *c*), the average of synaptic couplings (i.e. the charge associated to each postsynaptic potential; *Δ_EE_*, *Δ_EI_*, *Δ_IE_*, *Δ_II_*), and their standard deviations (i.e. *sΔ_EE_*, *sΔ_EI_*, *sΔ_IE_*, *sΔ_II_*):

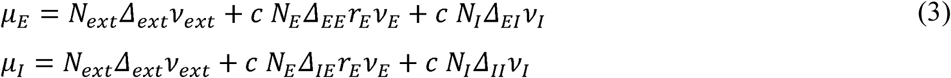

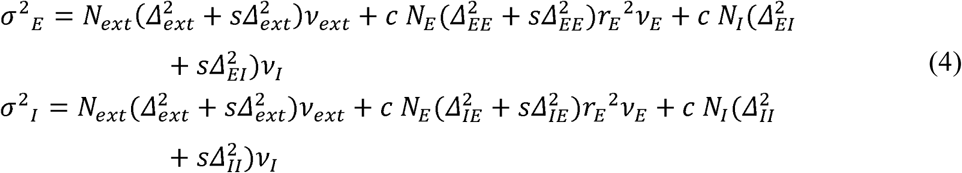

Following (Gigante et al. 2015), the dynamical filtering effects of AMPAr- and GABAr-mediated synapses were included by replacing the presynaptic mean firing rate *v* in eqs. 3-4 by their low-passed version 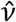:

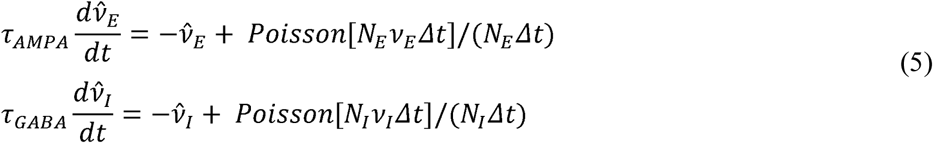

Equations 3-4 also included the effect of homosynaptic short-term synaptic depression at excitatory synapses by *r_E_*, evolving in time as

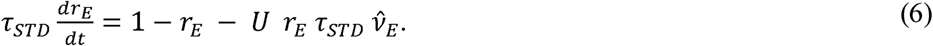

The simulation code together with a Jupyter notebook for its reproducible quantitative illustration have been made available on FigShare.com (DOI 10.6084/m9.figshare.6531293^1^).

## Results

To investigate circuit and cellular anomalies in an *in vitro* model of Fragile X syndrome, we prepared cortical cultures from Fmr1 knockout (KO) and wild-type (WT) control mice over seven distinct cell dissociation sessions, each leading to a series of cultures that we refer to as *sister* cultures. For monitoring and quantification of the electrical correlates of circuit dysfunction, we plated the cells on 62 substrate-integrated microelectrode arrays (MEAs; n = 37 for KO, n = 25 for WT) and cultured them for up to five weeks. We employed MEAs to non-invasively monitor extracellularly the coordinated electrical activity that spontaneously emerges *in vitro*. Presented are pooled data across *sister* cultures and culture sessions. Single-cell intracellular experiments were performed by patch-clamp recordings on a total of 100 neurons, plated on 100 glass coverslips (50 for each genotype), as we quantified the passive and active cell membrane electrical properties and the identity, amplitude, and frequency of spontaneous synaptic currents.

### Mature KO neuronal networks were spontaneously more active than controls

We followed over time the rate of spontaneous action potentials (i.e. the firing rate), detected across all microelectrodes of each MEA (Fig. 2A). We recorded the network activity at 7, 9, 14, 21, 28 and 35 days *in vitro* (DIVs) for at least 30 mins per MEA and day of experiment. We analyzed recordings obtained from a large number of MEAs, namely (KO) n = 29 for DIV7, n = 22 for DIV9, n = 25 for DIV14, n = 26 for DIV21, n = 9 for DIV28, and n = 10 for DIV35, and (WT) n = 25 for DIV7, n = 15 for DIV9, n = 25 for DIV14, n = 25 for DIV21, n = 25 for DIV28, and n = 18 for DIV35.

A well-known feature of cortical neuronal network development *ex vivo* is the spontaneous emergence of patterned electrical activity. Its evolution over time parallels the progression of network maturation, synaptogenesis, and pruning of synaptic connections (Kamioka et al. 1996; Marom and Shahaf 2002; Giugliano et al. 2004). Testing the hypothesis of whether the Fragile X network developed abnormal synaptic connections or displayed an altered maturation over time, we examined the rate of spontaneous action potentials over time. All networks displayed significant differences when their firing rate at early (i.e. DIV7-9) and late developmental stages (i.e. DIV14-35) were compared (Fig. 2A), confirming that both in KO and WT network development *ex vivo* occurs over a period of 2-3 weeks. However, while the firing rate in WT networks reached a peak at DIV21 and then decreased to a steady-state at DIV28-35, the firing rate in KO networks did not show any decrease after reaching its peak at DIV28 and it significantly exceeded the firing rate in WT from DIV21 onwards. This suggests that no refinement of the synaptic connections took place in KO, compared to WT.

More specifically, during the first two weeks *in vitro*, KO seemed less active than WT, but the difference was not significant (KO: 26 ± 3 at DIV7, 47 ± 5 at DIV9, 174 ± 18 spike/s at DIV14; WT: 42 ± 3 at DIV7, 70 ± 4 at DIV9, 197 ± 11 spike/s at DIV14). As time progressed, differences between genotypes became apparent: KO were less active at DIV21 than WT (KO: 167 ± 15 spike/s; WT: 208 ± 12 spike/s; p < 0.01) but later became more active (KO: 195 ± 17 at DIV28, 176 ± 12 spike/s at DIV35; WT: 121 ± 11 at DIV28, 111 ± 14 spike/s at DIV35; p < 0.01).

An in depth analysis of the data revealed that the spontaneous electrical activity was organized as irregular episodes of network-wide synchronized action potentials (APs) in both KO and WT networks (Fig. 1). We refer here to these episodes as network *bursts*, defined as cases in which the APs detected at distinct microelectrodes of the same MEAs show a certain degree of temporal overlap with each other’s (see the Methods section). Similarly to the firing rate, the *burst* rate is known to correlate with network development *ex vivo* (Ichikawa et al. 1993; Kamioka et al. 1996; Marom and Shahaf 2002). *Burst* rate displayed significant differences as early (i.e. DIV7-9) and late stages (i.e. DIV14-21) were compared (p < 0.01; Fig. 2B). Both KO and WT reached a peak in *burst* rate at DIV14, but KO networks maintained a higher rate of *bursting* at full maturity *in vitro* significantly exceeding the *burst* rate of WT. This suggest that network maturation was neither accelerated nor slowed down, but that mechanisms for the fine regulation of existing connections failed to intervene.

Specifically, during the first two weeks *in vitro* KO seemed less prone to synchrony than WT, although the differences were not significant (KO: 1.4 ± 0.1 at DIV7, 3 ± 0.3 at DIV9, 9 ± 0.8 burst/min at DIV14; WT: 1.9 ± 0.2 at DIV7, 4.4 ± 0.5 at DIV9, 9.4 ± 0.6 burst/min at DIV14). As time progressed, differences became significant: KO had lower *burst* rate at DIV21 than WT (KO: 5.8 ± 0.5 burst/min; WT: 7.05 ± 0.4 burst/min; p < 0.05) but had more than WT at DIV35 (KO: 5.5 ± 0.3 burst/min; WT: 3.6 ± 0.3 burst/min; p < 0.05). Fisher's least significant difference post-hoc test in the ANOVA was used in the analyses to assess significance: for Fig. 2A-B, the main effect for type (KO/WT) was not significant (p = 0.4), while the main effect for DIVs and the interaction between type and DIVs was significant (p < 0.05). Overall, these results indicate that the KO mature electrophysiological phenotype displayed signs of excitatory hyper connectivity, inhibitory hypoconnectivity, or cellular hyperexcitability.

### Quantitative differences in network *bursting* throughout *ex vivo* development

To extend our insight into the observed alterations, we next performed an extensive characterization of network *bursting* in KO and WT networks. We analyzed the distribution of *burst* duration, of number of APs per *burst*, and of inter-*burst* interval (Fig. 3). Consistent with their definitions, *burst* duration and the number of APs per *burst* shared similarities in their evolution over time: at full maturity *in vitro*, KO networks generated longer *bursts* (Fig. 3A), each composed of more numerous APs than WT (Fig. 3B).

**Figure 3.**
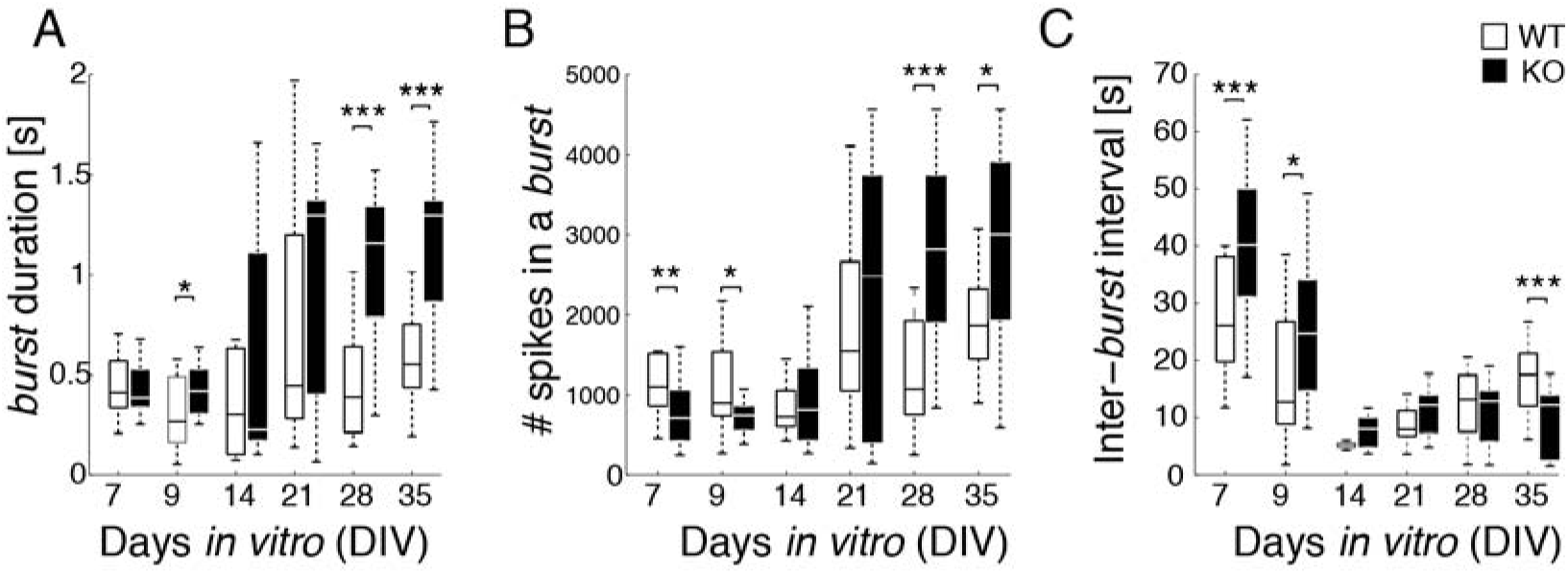
Detailed analyses of the network activity during development *ex vivo*. Box plots, displaying median, lower and upper quartiles (i.e. 25% and 75%) of the distributions, as well as lowest and highest sample values, are used to depict the features of network *bursting*. The distributions of the duration of each *burst*, of the number of spikes within the *burst*, and of the interval between one *burst* and the successive one are compared between KO (black bars) and WT networks (white bars). At full *ex vivo* maturity (>DIV28), KO networks display (**A**) significantly longer *bursts* (p < 0.001). Consistently, at full maturity KO *bursts* contain considerably more spikes than controls (p < 0.001 at DIV28 and p < 0.05 at DIV35) (**B**), but they contain less spikes than WT during network maturation (p < 0.01 at DIV7, p < 0.05 at DIV9). Extending the analysis of Fig. 2B, both KO and WT display (**C**) a progressive shortening of the inter-*burst* intervals over time. However, KO inter-*burst* intervals are significantly longer than in WT during network maturation (p < 0.001 at DIV7, p < 0.05 at DIV9). At full maturity, KO networks have significantly shorter inter-*burst* intervals than WT (p < 0.001 at DIV35).

Specifically, during the first two weeks *in vitro* the median values for *burst* duration for KO and WT networks (KO: 386 at DIV7, 421 at DIV9, 225 ms at DIV14; WT: 414 at DIV7, 266 at DIV9, 303 ms at DIV14) and for the number of APs per *burst* (KO: 710 at DIV7, 741 at DIV9, 812 at DIV14; WT: 1105 at DIV7, 899 at DIV9 725 at DIV 14) were not different. The distributions of the number of APs per *burst* were however significantly different at DIV7 (p < 0.01) and DIV9 (p < 0.05) in KO and WT, consistent with the low APs rates at the same age presented in Fig. 2A.

At full maturity *in vitro* (DIV28-35), both *burst* duration (p < 0.001) and the number of APs per *burst* (p< 0.001 at DIV28, p < 0.05 at DIV35) were significantly different between KO and WT (Fig. 3A-B), consistent with the higher APs rates in KO (Fig. 2A). Also, the median values were larger in KO than WT for *burst* durations (KO: 1297 at DIV21, 1156 at DIV28, 1297 ms at DIV 35; WT: 447 at DIV21, 390 at DIV28, 552 at DIV35) and for the number of APs per *burst* (KO: 2486 at DIV21, 2816 at DIV28, 3004 at DIV 35; WT: 1550 at DIV21, 1077 at DIV28, 1872 at DIV35).

The comparison of the distributions of inter-*burst* intervals in KO and WT (Fig. 3C) recapitulated the mean *burst* rate differences (Fig. 2B), with KO networks being more active than WT. Specifically, during the first two weeks *in vitro*, the median values for inter-*burst* duration were higher for KO than WT networks (KO: 40 at DIV7, 25 at DIV9, 8 s at DIV14; WT: 26 at DIV7, 13 at DIV9, 5 s at DIV14), and the distributions were significantly different at DIV7 (p < 0.001) and DIV9 (p < 0.05). At full maturity *in vitro* (DIV28-35), the distributions were significantly different (p < 0.001 at DIV35; Fig. 3C) with shorter median intervals in KO than in WT (KO: 12 at DIV21, 13 at DIV28, 12 s at DIV35; WT: 8 at DIV21, 13 at DIV28, 18 s at DIV35). Taken together, these data indicate that perturbed intrinsic cellular mechanisms, underlying the termination of each *burst*, could also play a role in KO networks.

### Qualitative differences in network *bursting* and intra-*burst* complexity

An extended statistical characterization of *burst* durations and inter-*burst* intervals, by directly estimating and comparing the cumulative distribution functions (cdf) and the probability distribution densities, is shown in Figure 4A-B. The figure displays both quantities over time and outlines the qualitative differences of *bursting* in KO networks and WT controls.

**Figure 4.**
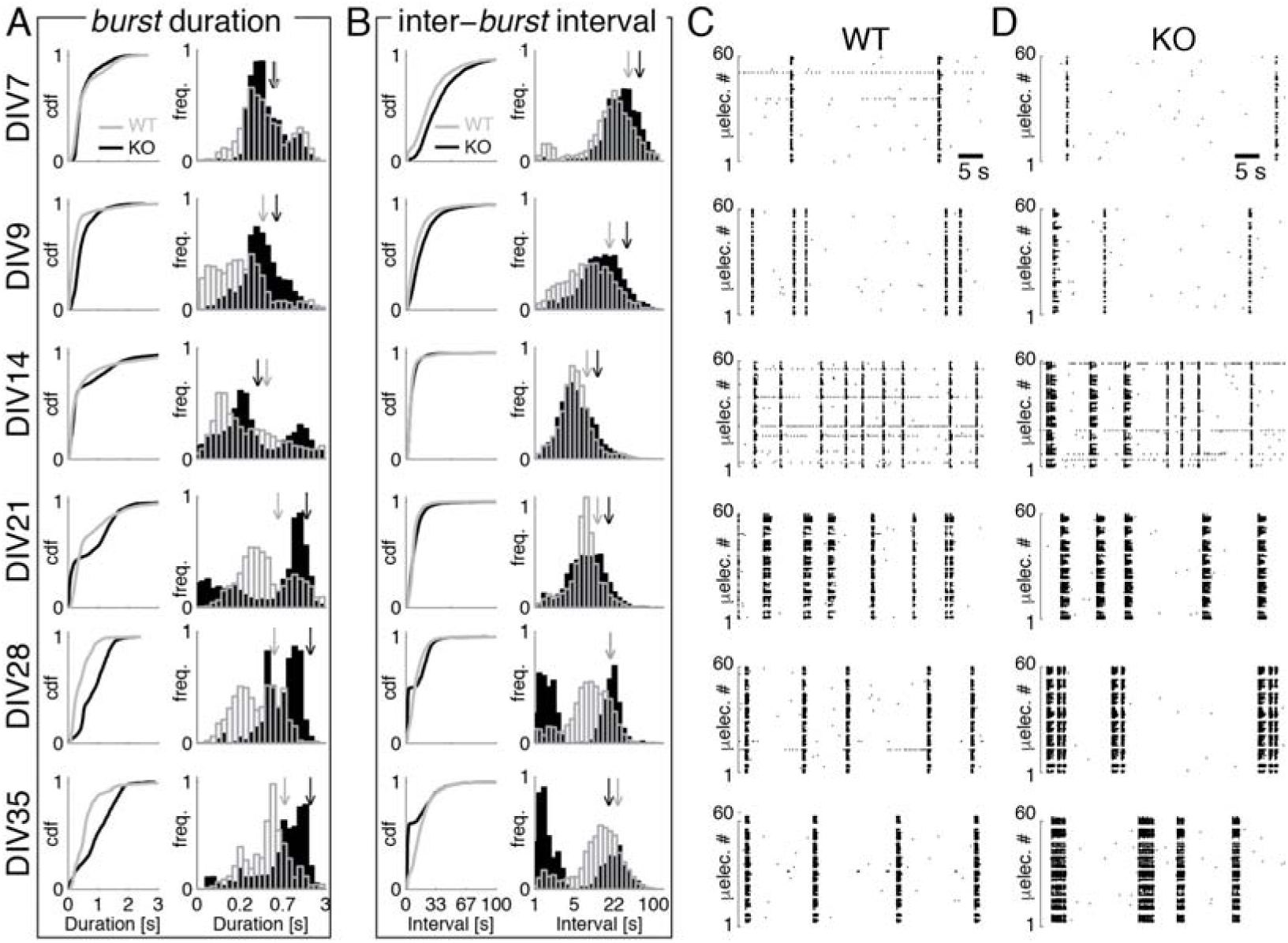
Evolution overview of the spontaneous activity during network development *ex vivo*. The analyses of Figs. 2-3 were further extended to visualize the cumulative distributions and probability densities for *burst* durations and inter-*burst* intervals, in KO (black lines and bars) and WT controls (grey lines and bars). Arrows indicate the median values; note the logarithmic horizontal scale. Over a period of 5 weeks *in vitro*, the analysis of *burst* durations reveals (**A**) the appearance of long *bursts* at DIV14 in KO networks in addition to short ones, as the distribution becomes bimodal. This is apparent as an additional peak in the probability density, which, from DIV21 onwards, dominates the distribution. The analysis of the inter-*burst* intervals (**B**) reveals the appearance of short intervals at DIV28, in addition to longer ones. This reflects the existence of a complex *burst* profile (Fig. 5) and it is apparent as an additional peak in the probability density, which remains bimodal until DIV35. Visual inspection of 50 s long sample spike-trains, detected across the MEAs and represented as raster plots in control (**C**) and KO networks (**D**), confirms that at full maturity (>DIV28) KO networks generate longer and more complex *bursts*.

With regards to *burst* duration, during the early stage of development *ex vivo* KO and WT networks initially generated similar *bursts*. However, starting from DIV14, the distribution of *burst* duration became bimodal in KO networks and skewed towards higher values (after DIV21), while remaining unimodal in WT. This indicates that both long and short *bursts* coexisted, and their longer *bursts* ultimately took over. Indeed, at DIV35 the distributions for KO and WT again turned into unimodal profiles, although they were still quantitatively different to KO networks generating longer *bursts*. For example, at 7DIV a randomly chosen *burst* from KO networks likely had a similar duration of a *burst* taken randomly from WT. Instead, by DIV28-35 the majority (>75%) of the *bursts* from KO networks were longer than a smaller fraction of *bursts* (<25%) from WT. This is apparent in the sample trains of APs (Fig. 4C-D) detected simultaneously across distinct microelectrodes of the MEAs and visualized as raster diagrams, over 50 s of recordings.

When we examined the inter-*burst* interval distribution, we found that the opposite sequence of events took place: until DIV21, KO and WT networks generated *bursts* with unimodal distributions and with KO networks first generating *bursts* less frequently than in WT (DIV7-14) and then with comparable frequency (DIV21). However, from DIV28 onwards, KO networks generated a bimodal distribution of inter-*burst* intervals, with WT networks only having a weak trend to perform the same. This is apparent in the raster plots (Fig. 4C-D), where successive *bursts* in KO were separated alternatively by short or long pauses. Indeed, only in KO networks short inter-*burst* intervals were equally frequent than the longer intervals.

Taken together, these results indicate that a dysfunction of the intrinsic mechanisms underlying *burst* termination is unlikely, and that network excitability as sustained by excitatory and inhibitory connectivity may be occurring. The emerging picture is that episodic neuronal synchronization in KO networks was associated with more complex phenomena than in WT. For this reason, we examined in greater details the time course underlying the instantaneous rate of APs, during each network synchronization (Fig. 5). We found that each *burst* could be distinguished in an early and a late phase, which were previously attributed to intrinsic and synaptic mechanisms, respectively (Pulizzi et al. 2016). The early phase was characterized by a sudden, exponential, increase over time of the instantaneous firing rate, in both KO and WT networks (not shown). At DIV35, we found substantial qualitative differences in the late phase of the *bursts*, persisting for KO networks for several seconds (Fig. 5C-D). As the power spectrum of the late phase was estimated (Fig. 5E-H), we found in KO but not in WT a prominent intra-*burst* oscillation of the instantaneous firing rate. Starting several hundreds of milliseconds from the burst profile peak amplitude, such oscillatory activity was clearly dominated by *beta*-band power (~18 cycle/s; in 8 out of 10 MEAs) and it was completely absent in WT (10 out of 10 control MEAs). These results suggest that in KO, the synaptic mechanisms underlying network excitability are significantly altered compared to WT, with a new dynamical regime emerging during the late component of the *bursts*.

**Figure 5.**
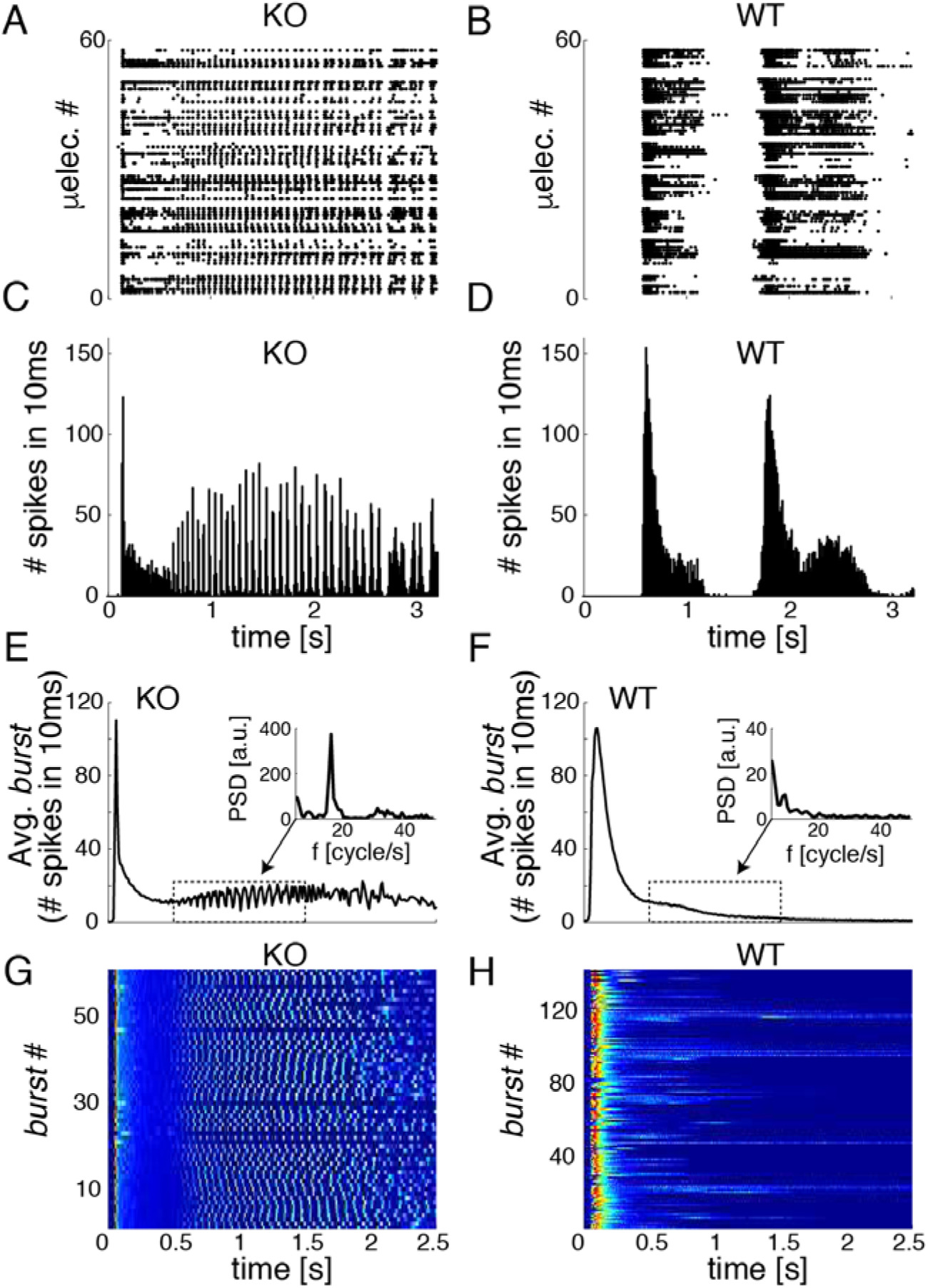
*Burst* complexity reveals oscillations in the *beta* frequency band at DIV35. Sample spontaneous *bursts* in KO networks (**A**) and WT controls (**B**) are represented as a raster diagram and show substantial differences in the temporal progression of APs activity and its synchronization. When the spike-time histograms are estimated for the same data in KO networks (**C**) and controls (**D**), a prominent intra-*burst* modulation of the rate of firing in KO becomes apparent. This is even more convincing when all the *bursts* detected during the same recording session are averaged together (**E**-**F**), and the power-spectral density is computed over the *burst*’s tail (indicated by the dashed box): *beta* frequency range oscillations occur in KO networks but are absent in WT. This is further visually rendered (**G**-**H**) with pseudo-colours, displaying the intra-*burst* firing rate profile across several individual *bursts*.

### Altered single-neuron passive properties and excitability

To rule out the hypothesis that major single-cell intrinsic alterations in KO were responsible for the network-level abnormalities observed, we performed whole-cell current-clamp recordings from the soma of KO and WT cultured neurons. In these experiments, the cellular phenotype can be separated from the network complexity, as a single cell at that time is under the control and observation of the experimenter. We then first examined the passive electrical properties of the cells, by determining their membrane time constant. This was quantified by standard methods, upon fitting an exponential function to the membrane potential trajectory, as it recovered to the resting membrane potential, after a brief hyperpolarizing current pulse (Fig. 6A). We found no significant differences between KO and WT neurons, during early stages of development (DIV7-14). However, at DIV21-28, KO neurons were significantly slower in recovering to their resting membrane potentials (p < 0.05 at DIV21 and p < 0.001 at DIV28; n = 10 for KO and n = 10 for WT). These data suggest that the integrative properties of the neuronal membrane in KO are less prominent than in WT, which is at odds with the hypothesis of intrinsic cellular hyperexcitability.

**Figure 6.**
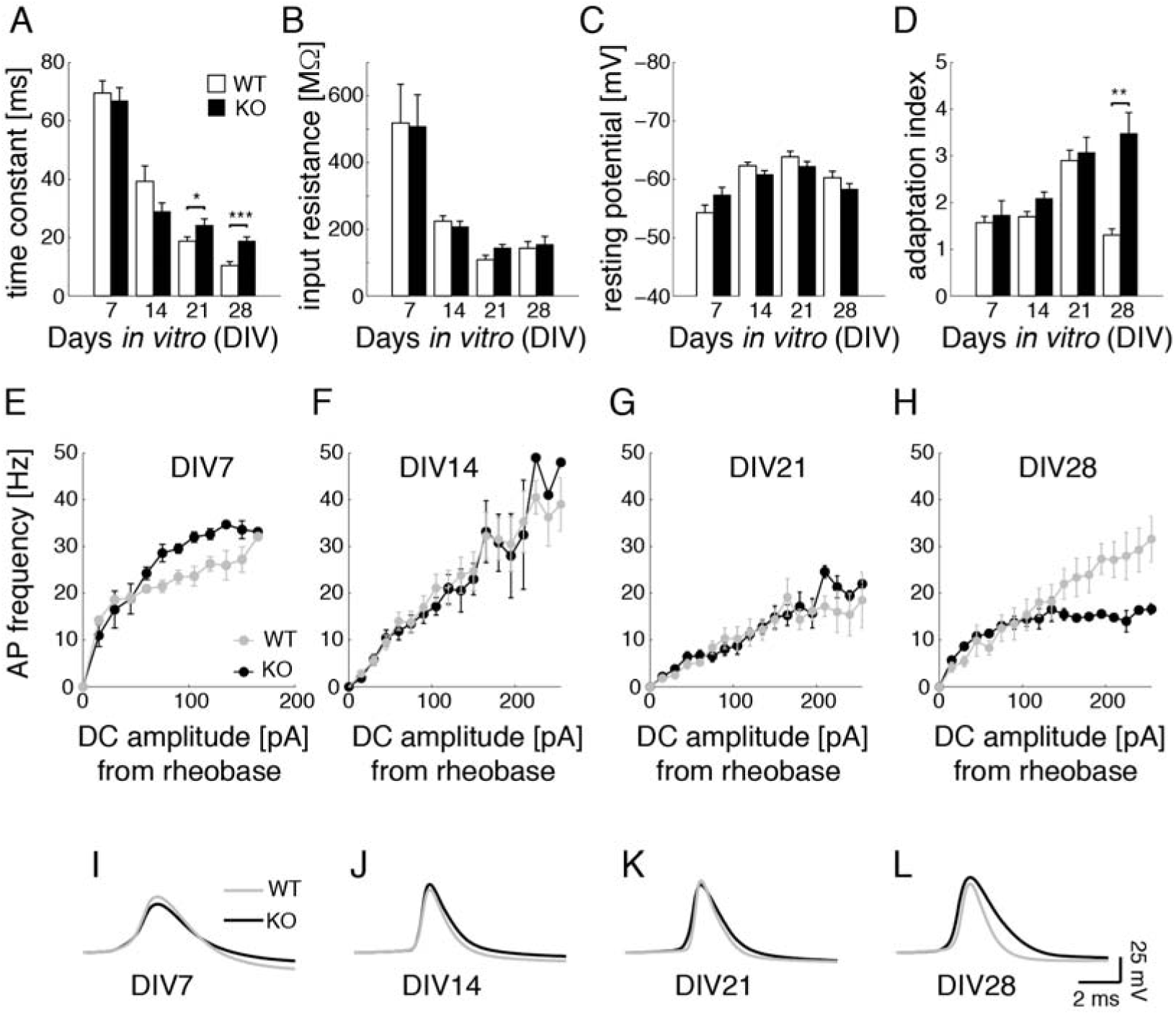
Single-cell passive and active properties during development *ex vivo*. Patch-clamping was employed to probe single-cell passive and active membrane properties, over n = 10 cells per condition and per DIV. Data are shown as mean ± SEM and significance is assessed by a two-sample t-test. Average membrane time constant (**A**), input resistance (**B**), and resting potential (**C**) are quantified by standard protocols in KO (black bars) and WT neurons (white bars). Concerning the passive properties, KO and WT neurons are indistinguishable with the exception of a significantly longer membrane time constant at full maturity (>DIV21). Concerning excitable properties, the spike-frequency adaptation index (**D**) and the APs frequency *versus* current intensity (f-I) curves (**E**-**H**) are measured (KO: black markers and lines; WT: grey markers and lines): KO and WT neurons are indistinguishable with the exception of a stronger adaptation index and consequently of a less steep f-I curve, at full maturity (>DIV28). No differences are observed in the average AP shapes (**I-L**), with the exception of a less pronounced AP repolarization in KO neurons at full maturity (KO: black markers and lines; WT: grey markers and lines).

We found no differences between KO and WT neurons in terms of their apparent input resistance (Fig. 6B) and resting membrane potential (Fig. 6C), which were in physiological ranges throughout the development *ex vivo*. These results then further support that KO cells are not more excitable than WT.

To investigate the intrinsic excitability directly, we examined the generation of a train of action potentials in KO and WT, studying the ability of individual cells to respond to the injection of depolarizing DC current pulses. No significant differences were found in the values of the rheobase current (not shown), which is the minimal current amplitude to elicit regular AP firing, with the exception of the values at DIV21 (KO: 107 ± 10 pA; WT: 160 ± 15 pA; p < 0.05). The AP responses were first quantified in terms of the spike-frequency adaptation index (Fig. 6D), i.e. upon dividing the duration of the last inter-spike interval of the train by the duration of the first. At DIV28, KO neurons had a significantly larger adaptation index (p < 0.01; n = 8 for KO and n = 8 for WT), implying an overall reduced propensity to fire APs at high frequency. This was confirmed by investigating the AP frequency *versus* current amplitude curves (Fig. 6E-H), where at DIV28 a much slower slope of the curve was apparent in KO than WT neurons, while sharing similar rheobase currents (KO: 92 ± 11 pA; WT: 160 ± 34 pA; p = 0.14).

Finally, we examined the average AP shape during maturation and found that it was not significantly different until DIV21. Indeed, at DIV28 a slower repolarization characterised the AP shape in KO neurons than WT, representing a hallmark for a weaker reset of the membrane potential during repetitive AP firing. This observation is consistent with the less steep AP frequency-current curve (Fig. 6H). These results indicate that KO cells are not more excitable than WT, suggesting that synaptic and not intrinsic cell properties underlie the network abnormalities.

### Quantitative differences in synaptic currents underlying *bursting*

We then hypothesised that the observed alterations (Fig. 2-4) were associated with an unbalance of excitatory *versus* inhibitory synaptic transmission. To directly test this hypothesis, we performed voltage-clamp experiments and electrically isolated spontaneous excitatory (n = 8 for KO and n = 8 for WT, for each DIV) and inhibitory currents (n = 8 for KO and n = 8 for WT, for each DIV) (Fig. 7C,G), upon matching by the holding potential the Nernst equilibrium potential of AMPA/NMDA receptors and or GABAA receptors, respectively. This makes it possible to electrically “cancel” alternatively the excitatory or inhibitory components of spontaneous synaptic transmission.

**Figure 7.**
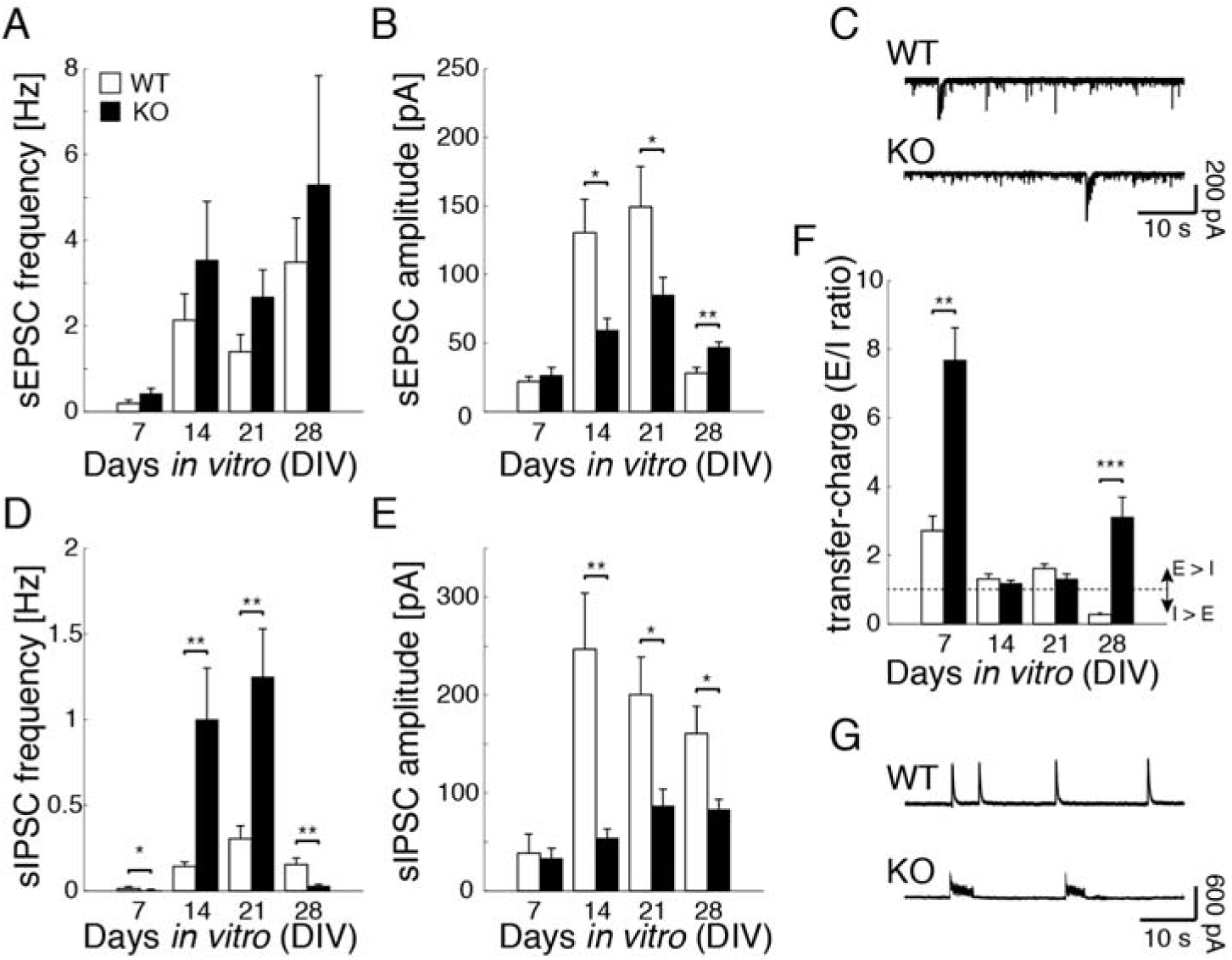
Analyses of synaptic currents during development *ex vivo*. Voltage-clamp recordings of spontaneous postsynaptic excitatory (inhibitory) synaptic currents (sEPC, sIPSC) were performed, holding neurons at −70 mV (0 mV), in KO (black bars) and in WT (white bars), over n = 8 cells per condition and per DIV. Data are shown as mean ± SEM. The developmental progression of Figs. 2B-3C is reflected in the mean frequency of sEPSC (**A**) for both KO and WT, without differences. In KO, sEPSC amplitudes are (DIV14-21) significantly smaller during maturation (DIV14**-**21) but, upon reaching full maturity (DIV28), they are larger than control. sEPSCs representative examples are shown (**C**) for KO and WT neurons, at DIV14. Inhibitory synaptic currents differ significantly in KO compared to WT (**D**-**E**). While sIPSC amplitudes are smaller in KO than in WT throughout development *ex vivo* (p < 0.01 at DIV14 and p < 0.05 at DIV21-28), sIPSC frequency is larger during network maturation (p < 0.01 at DIV14-21) and smaller than in WT upon reaching full maturity (p < 0.01 at DIV28). The balance between excitation and inhibition in KO neurons is significantly shifted towards excitation than in WT, in terms of electrical charge transferred and quantified by the ratio of the areas below sEPSC and sIPSC recorded waveforms. sIPSCs representative examples are shown (**G**) for KO and WT neurons, at DIV14.

When the frequency of spontaneous excitatory and inhibitory postsynaptic currents (sEPSC and sIPSC) was studied (Fig. 7A,D), we observed that KO neurons received on average more sEPSC per unit of time than WT (KO: 0.42 ± 0.12 at DIV7, 3.54 ± 1.28 at DIV14; 2.67 ± 0.59 at DIV21, 5.3 ± 2.55 Hz at DIV28; WT: 0.19 ± 0.08 at DIV7, 2.13 ± 0.61 at DIV14, 1.4 ± 0.41 at DIV21, 3.49 ± 1.03 Hz at DIV28; Fig. 7A), although these differences were not significant. Instead, for sIPSC the difference between KO and WT neurons was highly significant (p < 0.05 at DIV7, p < 0.01 at DIV14-28). Specifically, during early stages of development *ex vivo*, KO neurons received significantly more sIPSC per unit of time than WT, while at DIV28 significantly less sIPSC than WT (KO: 0.001 ± 0.0 at DIV7, 1. ± 0.28 at DIV14, 1.25 ± 0.26 at DIV21, 0.03 ± 0.01 Hz at DIV28; WT: 0.02 ± 0.01 at DIV7, 0.14 ± 0.02 at DIV14, 0.30 ± 0.08 at DIV21, 0.15 ± 0.04 Hz at DIV28; Fig. 7D).

While the frequency of sEPSC and sIPSC is highly correlated with network-wide *bursts* (Fig. 2-4), their mean amplitude is proportional to the efficacy of synaptic transmission (Fig. 7B,E). We then found that during early stages of *in vitro* maturation, sEPSC amplitudes were on average significantly weaker in KO neurons than WT (p < 0.01, DIV14-21). Instead, at DIV28 we observed the opposite phenomenon, as KO neurons received stronger sEPSC than WT controls (KO: 26±6 at DIV7, 59 ± 8 at DIV14, 85 ± 12 at DIV21, 47 ± 4 pA at DIV28; WT: 22 ± 3 at DIV7, 131 ± 23 at DIV14, 149 ± 30 at DIV21, 28 ± 4 pA at DIV28; Fig. 7B).

Instead, sIPSC amplitudes were always significantly weaker (p < 0.01 at DIV14 and p < 0.05 at DIV21-28) in KO neurons than WT (KO: 33 ± 10 at DIV7, 54 ± 9 at DIV14, 87 ± 17 at DIV21, 83 ± 10 pA at DIV28; WT: 39 ± 20 at DIV7, 247 ± 57 at DIV14, 200 ± 38 at DIV21, 161 ± 27 pA at DIV28; Fig. 7E).

Finally, the charge transfer associated with sEPSC and sEPSC could be estimated by the area below their trajectories. Figure 7F summarises our findings, showing significant differences at DIV7 (p < 0.01) and DIV28 (p < 0.001) between KO neurons and WT controls. Specifically, in KO neurons the total charge transferred by excitatory synapses was on average always larger than the one transferred by inhibitory synapses, while for WT neurons inhibition prevailed at DIV28 (KO: 7.68 ± 2.31 at DIV7, 1.18 ± 0.25 at DIV14, 1.31 ± 0.41 at DIV21, 3.12 ± 1.42 at DIV28; WT: 2.72 ± 0.86 at DIV7, 1.31 ± 0.43 at DIV14, 1.62 ± 0.37 at DIV21, 0.28 ± 0.09 at DIV28). Taken together, these results indicate that KO networks are characterised by more numerous or more effective excitatory synaptic connections and by less numerous or less effective inhibitory synaptic connections than in WT.

### A minimal mathematical model reproduces network *bursting* alterations

In a synthesis effort, we designed and simulated a mathematical model of neuronal networks (Fig. 8). We aimed to explain the observed electrophysiological phenotypes in KO and WT, in the simplest qualitatively terms and upon linking synaptic properties to network-level observables. The model considers the interplay between two distinct mechanisms: a positive feedback, arising from recurrent excitatory connections, and a negative feedback, related to cellular and synaptic fatigue. From this model, we found that such an interplay is responsible for the irregular *bursting* (Fig. 8B-C), which occur as a robust emerging phenomenon without fine tuning of the model parameters. Interestingly, as soon as the inhibitory connections were downregulated in number and efficacy, the inter-*burst* intervals decreased, *bursting* became more regular, and the duration of each *burst* increased, as shown in Figs. 2-4. When excitatory connections were further increased (Fig. 8D-E), each *burst* became characterised by prominent oscillatory activity in its late phase, qualitatively similar to the experiments of Fig. 5.

**Figure 8.**
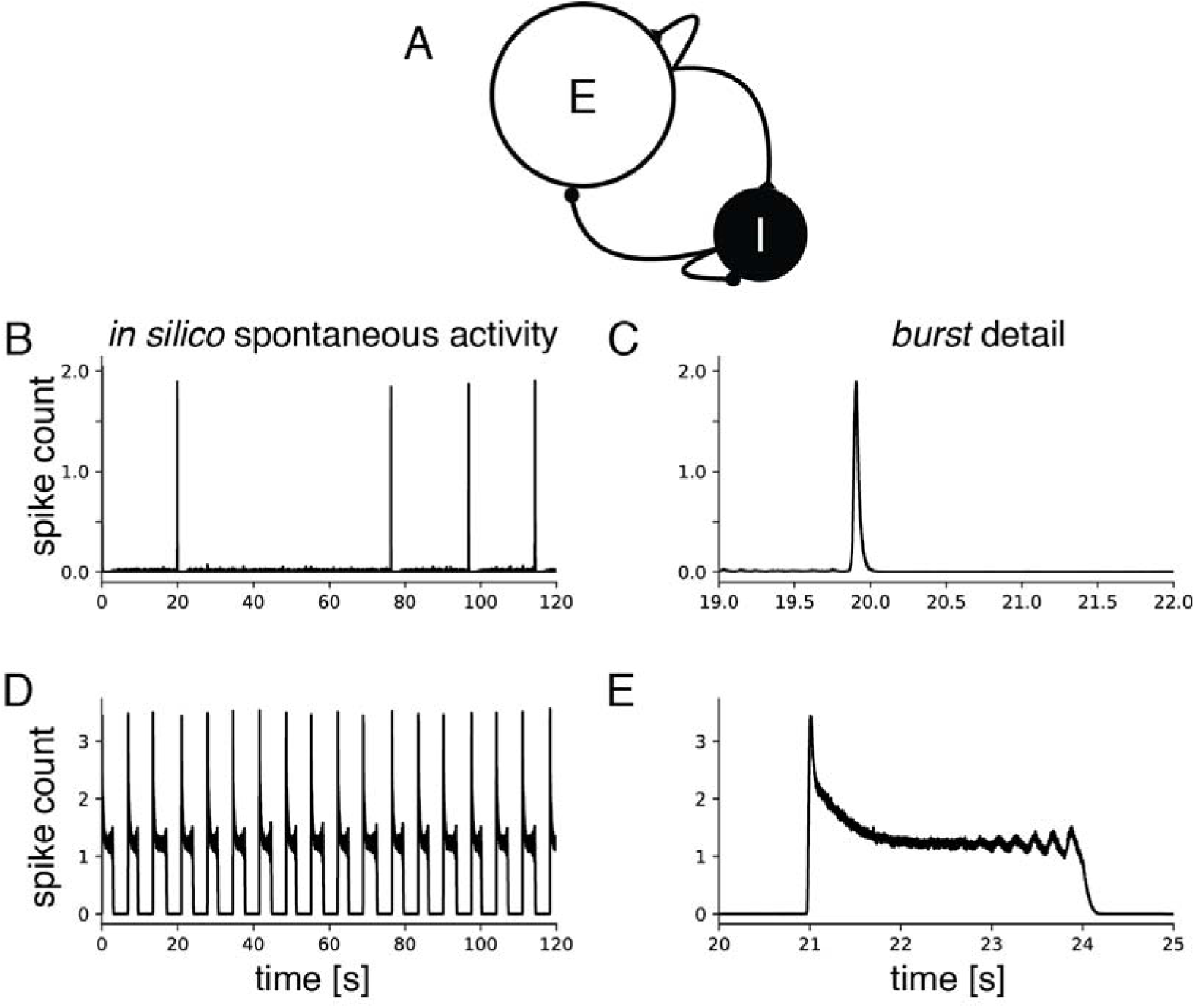
A minimal explanatory model qualitatively reproduces our experimental observations. A firing-rate mathematical model was simulated to explore *in silico* the links between synaptic connectivity and network-wide electrical phenomena. The model features two neuronal subpopulations of excitatory and inhibitory neurons, identified as “E” and “I” in the sketch (A), connected reciprocally and recurrently by chemical synapses. Computer simulations illustrate that irregular *bursting* occurs spontaneously (B-C), emerging from the interplay between positive and negative feedback mechanisms. Mimicking the synaptic dysfunctions in KO, we downregulated inhibitory synaptic connections and upregulated excitatory connections, while keeping all the other numerical parameters identical. Such changes were sufficient (D-E) to decrease inter-*burst* intervals, increase *burst* durations, and reveal oscillations in the late phase of the *burst*, as in the experiments.

These results indicate that an alteration in the balance of excitatory-inhibitory connection efficacies is sufficient to explain the network-level electrical disturbances observed in KO.

## Discussion

We studied an *in vitro* model of Fragile X syndrome, suitable for longitudinal investigation of network properties and for cellular and synaptic detailed analyses.

We first observed a clear alteration in the spontaneous firing in KO networks compared to WT, which displayed progression towards a mature activity pattern through time. Prior to DIV5, isolated APs were fired *asynchronously* across the culture. Later, regular occurrence of *bursts* of synchronised APs emerged and evolved, over the following 2-3 weeks, into a more complex pattern (Kamioka et al. 1996; Wagenaar et al. 2006; Golshani et al. 2009). As spontaneous electrical activity mirrors cellular maturation and synaptogenesis (Ichikawa et al. 1993; Kamioka et al. 1996; Marom and Shahaf 2002; Giugliano et al. 2004), we monitored the activity of our networks by MEA, and we obtained a direct insight into the maturation of the synaptic connectivity.

Fmr1 KO network*s* displayed shorter, sparser and rarer bursts at early development and longer, denser and more frequent ones at the matured stage, compared to WT. Despite obvious differences from *in vivo* intact cortical activity, our findings reconcile the observations of (Hays et al. 2011) and (Gonçalves et al. 2013) of prolonged cortical “up states” observed in Fmr1 KO mice only after 3 weeks of age and not earlier, as our *bursting* may be considered a simpler correlate of the activity in the intact cortex. In our experiments, significant alterations of network *bursting* arose after 3 weeks *in vitro* and were detected at DIV28.

Our main observation is that Fmr1 KO networks were hyperactive, in line with previous reports (Gonçalves et al. 2013; Rotschafer and Razak 2013; Scharkowski et al. 2018). These alterations in network activity occurred mostly after maturity *in vitro and* were associated with non-trivial electrophysiological characteristics. These could not be attributed to alterations in intrinsic cellular excitability, as single-cell experiments did not reveal any differences contributing to enhancing cellular responsiveness to external inputs. Therefore, the observed abnormalities arose from alterations in synaptic transmission and in synaptic connectivity. We previously reported hyper-connectivity of excitatory neurons in the developing intact cortex of Fmr1 KO mice and that this feature was spontaneously reversed to physiological levels after 4-5 weeks of development (Testa-Silva et al. 2012). In addition, we note that a higher cortical dendritic spine density had been observed in Fmr1 KO adult mice (McKinney et al. 2005) and a delay of spine maturation in FXS was also reported (Cruz-Martin et al. 2010). Thus, we conclude that a hyper connectivity of glutamatergic connections is likely to characterise our *in vitro* circuits. We cannot rule out that by prolonging our experiments in time, we would also have observed a reversal of the hyperactivity, although the health and the long-term survival of cell cultures could then act as a confounding factor. However, even if Fig. 2A apparently indicates a delayed maturation in KO compared to WT, the rate of network *bursts* did not show the same delay (Fig. 2B). *Bursting* rather than APs firing is a better correlate of network maturation (Kamioka et al. 1996; Marom and Shahaf 2002). For this reason, we interpret our results as a developmental failure to downregulate the network excitability into physiological regimes, rather than a delayed connectivity maturation as *in vivo*. As in FXS both (sub)cellular mechanisms are likely to be disrupted, intervention strategies to correct them all might benefit from the advantages offered by our networks plated on MEAs, aiming at ultimately rescuing a physiological phenotype by pharmacology.

In our single-cell experiments, we did not observe significant functional differences in the excitatory transmission after the first week of development (Fig. 7A-B), despite reports on spine length and density abnormalities (Nimchinsky et al. 2001) and in AMPA/NMDA ratio differences (Harlow et al. 2010) during early cortical synaptogenesis. Besides an overexpression of excitatory currents, we also attribute synaptic abnormalities to a downregulation of GABAergic transmission, given the disruption of the excitation-inhibition balance and the substantial disruption in frequency and amplitude of spontaneous inhibitory postsynaptic currents (Fig. 7D-G). In fact, inhibitory transmission was dramatically altered, breaking down at DIV28 and thus was directly responsible for altering recurrent activity and transforming it into an intense hyper active regime (Chagnac-Amitai and Connors 1989).

As GABA is crucial for proper network maturation and wiring (Akerman and Cline 2006; Wang and Kriegstein 2008), deficits in GABAergic signalling may explain the lack of excitatory activity detected at DIV14-21. At that time, inhibitory events in KO networks had decreased amplitude and increased spontaneous frequency compared to WT. All these findings might not only indicate AMPA receptor internalization (Nakamoto et al. 2007), but also a reduced amount of GABAA receptors or their lower sensitivity. These could indeed be the direct result of an under expression of multiple subunits of the GABAA receptors (D’Hulst et al. 2006; Braat et al. 2015), which in WT acted as a dynamic negative feedback for the *burst* termination.

With regards to the overall network wiring, as for *in vivo* cortical “up states” *in vitro bursting* is also the direct result of recurrent synaptic connectivity (Marom and Shahaf 2002; Giugliano et al. 2004). Our findings on the disproportional spontaneous activity in Fmr1 KO networks are unlikely to be caused by an excess activation of the group I glutamate metabotropic receptors (mGluR5) (Hays et al. 2011) as the mathematical model replicated the complexity in *burst* shape upon alteration of fast (i.e. AMPA/GABA) not slow synaptic coupling mechanisms. Besides the efficacy of excitatory and inhibitory synaptic transmission, intrinsic cell excitability could have played a role in determining KO network activity, e.g., affected by a reduced expression of Kv4.2 potassium voltage-gated ionic currents (Gross et al. 2011). However, while such reduced expression was likely the cause of the differences in the AP shape at DIV28 (Kim et al. 2005) (Fig. 6L), we did not consider it as a major contributor to network hyperactivity in our experiments, as the APs frequency *versus* current intensity f-I curve of KO neurons displayed a less steep profile than in WT (Fig. 6H), and not vice versa. This might be the result of the cumulative inactivation of sodium currents (Fleidervish et al. 1996) whose persisting effect decreased excitability and it is not removed by strong hyperpolarizing potassium currents (Pampaloni et al. 2018) in KO compared to WT. Alternatively, the significantly lower slope of the APs frequency *versus* current intensity curves reported here at DIV28 (Fig. 6H) might be the result of a homeostatic attempt to compensate for the disruption of the excitatory-inhibitory balance (Fig. 7F) and of putative hyper-connectivity. Indeed, the balance between voltage-gated depolarizing and repolarizing ionic currents (e.g. sodium fast-inactivating currents, calcium-currents, delayed rectifier currents, etc.) regulate intrinsic cellular excitability and is also a known regulator of network activity (Turrigiano 2011).

The clear findings of oscillatory intra-*burst* activity in the *beta*-power frequency range in KO but not WT networks were unexpected. Previous mathematical work (Pulizzi et al. 2016) suggests that the interplay between excitation and inhibition might be responsible for these oscillations. An altered balance between excitation and inhibition could therefore have manifested itself in the oscillations, which are interestingly reminiscent of an *in vivo* (EEG) biomarker of another autism spectrum disorder syndrome with aberrant GABAA activity (Frohlich et al. 2016). However, an alternative hypothesis for this oscillatory activity may come from the known dysregulation of short-term synaptic plasticity in Fragile X (Testa-Silva et al. 2012). Indeed, computer simulations (Masquelier and Deco 2013) support the emergence of a complex *bursting* activity profile from the interaction of time-scales of distinct adaptation mechanisms, including short-term synaptic depression and facilitation. In our model, we could qualitatively replicate those slow intra-*burst* oscillations in the absence of any inhibition, suggesting an interplay between glutamatergic synaptic transmission and the negative feedback mechanisms underlying burst termination. In the model, this was explained however only as a consequence of a decrease in the number and strength of inhibitory synaptic connections and of an increase in the number and strength of the excitatory synaptic connections.

## Conclusions

Our study demonstrates that loss of Fmr1 in neurons results in impairments at multiple levels of circuit organization, largely involving both excitatory and inhibitory transmission and to a much lesser extent single-cell excitability. These alterations manifested themselves in complex forms at the network level, indicating that any therapeutic intervention might require timely and precise pharmacological modulation of synaptogenesis and of synaptic transmission during network formation in order to recover the WT phenotype *in vitro*. The experimental model of a disease “in a dish” that we employed, despite its obvious limitations in comparison to *in vivo* electrophysiology and molecular biology, might easily allow extensive screening of therapeutic strategies while accessing an easy to monitor quantitative observable such as the electrical phenotype of the network as a whole.

## Acknowledgments

We are grateful to Drs. S. Braat and J. Couto and to Mrs M. Malezadeh for discussions and assistance with animal breeding, and to Mr. D. Van Dyck and M. Wijnants for excellent technical assistance. Financial support from the European Union’s Horizon 2020 Framework Programme for Research and Innovation under the Specific Grant Agreement n. 785907 (Human Brain Project SGA2), the Belgian Science Policy Office (grant n. IUAP-VII/20), the Flemish Research Foundation (grant n. G0F1517N), the FRAXA research foundation is kindly acknowledged. The funders had no role in study design, data collection and analysis, decision to publish, or preparation of the manuscript.

## Contributions

AM, RFK, and MG designed and supervised the research. AM performed and analyzed the electrophysiological experiments. MG conceived the mathematical model and performed the numerical simulations. AM, RFK, and MG wrote the paper. All authors read and approved the final manuscript.

## Data accessibility statement

Authors confirm that the data underlying their findings are fully available. Relevant data sets and analysis scripts have been stored at FigShare.com (DOI 10.6084/m9.figshare.6531293^2^), including an index of the deposited data.

## Footnotes

1 Reviewers and Editors private access link: https://figshare.com/s/2558eab9d4ec9153beda

2 Reviewers and Editors private access link: https://figshare.com/s/2558eab9d4ec9153beda

